# Agency rescues competition for credit assignment among predictive cues from adverse learning conditions

**DOI:** 10.1101/2021.02.24.432808

**Authors:** Mihwa Kang, Ingrid Reverte, Stephen Volz, Keith Kaufman, Salvatore Fevola, Anna Matarazzo, Fahd H. Alhazmi, Inmaculada Marquez, Mihaela D. Iordanova, Guillem R. Esber

## Abstract

A fundamental assumption of learning theories is that the credit assigned to predictive cues is not simply determined by their probability of reinforcement, but by their ability to compete with other cues present during learning. This assumption has guided behavioral and neural science research for decades, and tremendous empirical and theoretical advances have been made identifying the mechanisms of cue competition. However, when learning conditions are not optimal (e.g., when training is massed), credit assignment is no longer competitive. This is a catastrophic failure of the learning system that exposes the individual’s vulnerability to form spurious associations in the real world. Here, we uncover that cue competition can be rescued when conditions are suboptimal provided that the individual has agency over the learning experience. Our findings reveal a new connection between agency over learning and credit assignment to cues, and open new avenues of investigation into the underlying mechanisms.

---

The ability to predict outcomes of biological significance is essential to fitness and survival. To do so, animals, including humans, must identify which stimuli among the many present provide relevant predictive information (Mackintosh, 1974; Dickinson, 1980); that is, they must solve the problem of structural credit assignment (Sutton & Barto, 2018). A dramatic example of this type of problem is provided by the COVID-19 pandemic, as societies around the world scrambled to ascertain which stimuli posed a risk of contagion and which did not. Failure to assign predictive credit to relevant stimuli (e.g., close physical contact with other individuals, shared meals, enclosed spaces) has led to dire consequences (e.g., Fisher et al., 2020), while misassigning credit to irrelevant stimuli (e.g., 5G mobile networks, mosquitoes, bleach) has likewise fostered maladaptive behaviors (Nowak et al., 2020; Krause et al., 2020). Given the intimate relationship between credit assignment and decision making, it is essential to elucidate how credit is apportioned among cues under various learning conditions.

Credit assignment is widely regarded as a competitive process in which the best predictor of the outcome acquires substantial credit over the course of learning at the expense of other predictors (e.g., Honey et al., 2020; Sutton & Barto, 1990; Delamater, 2012; Harris, 2006; Wagner 2003; McLaren & Mackintosh, 2000; Brandon et al., 2000; Pearce, 1994; Miller & Matzel, 1988; Wagner, 1981; Pearce & Hall, 1980; Mackintosh, 1975; Rescorla & Wagner, 1972). In support of this notion is evidence that cues compete for credit in a range of tasks (Kamin, 1968; Pavlov, 1927, Wagner et al., 1968, Rescorla, 1968, 1970) and species, from C. elegans to humans (Merritt et al., 2019, Prados et al., 2013, Blaser et al., 2006, Pearce et al., 2012, Kamin, 1968, Wagner et al., 1968, Beauchamp et al., 1991, Tobler et al., 2006; Dickinson et al., 1984). However, it has also long been known that cue competition is not ubiquitous (e.g., Durlach & Rescorla, 1980; Batsell & Batson, 1999; Vadillo & Matute, 2010; Maes et al., 2016) and can be disrupted across multiple learning conditions (reviewed in: Urcelay, 2017; Witnauer et al., 2014; Wheeler & Miller, 2008). One such condition is experiencing the trials in massed fashion (in rats: Stout et al., 2003; Sissons & Miller, 2009; in pigeons: Packheiser et al., 2020; in humans: Beesley & Shanks, 2012). This finding has profound implications because in our ever-complex world we are routinely bombarded with information presented in close succession. Diminished cue competition in such situations implies that incidental stimuli may highjack the learning system and wrongfully gain control over behavior.

Recently, Reverte et al. (2020) reported that granting rats agency over trial presentations protects cue-reward learning from the well-known deleterious effects of massed training (e.g., Lattal, 1999; Barela, 1999; Holland, 2000; Sunsay & Bouton, 2008). Here, we sought to determine whether agency over learning could specifically rescue competitive credit assignment under massed trial conditions. To this end, we embedded a trial self-initiating procedure within a powerful master-yoked design that allowed us to vary the degree of agency over learning while keeping the exposure to cue-reward relationships identical (Reverte et al., 2020). Using a variety of well-established and novel cue competition tasks, we found robust evidence of cue competition only in animals that had agency over learning. Importantly, this effect was not the consequence of differential levels of engagement, general discrimination ability or propensity to process compounded stimuli concurrently. Our data provide the first demonstration of a critical role for agency in credit assignment and open up new lines of neural and theoretical inquiry.

## Results

### Agency rescues the blocking effect from the deleterious effects of massed training

We first set out to test whether agency over learning rescues the blocking effect (Kamin, 1968) from the deleterious effects of massed training in (Fig. 1B). This and the remainder of the studies employed a within-subject design embedded within a between-subject master-yoked procedure (Fig. 1A). Master rats (Group Agency) were allowed to self-initiate their trials by performing a nose poke into a nose port at any point during a period of trial availability (max = 20 s) signaled by a nose port light (Reverte et al., 2020). On any given trial, a nose poke would turn on one or more 10-s cues of the visual and auditory modality. In contrast, yoked rats (Group Passive) received an identical sequence of events to their master counterparts (including the trial availability cue), but trial presentations were noncontingent on their behavior (i.e., standard Pavlovian conditioning).

**Figure 1.**
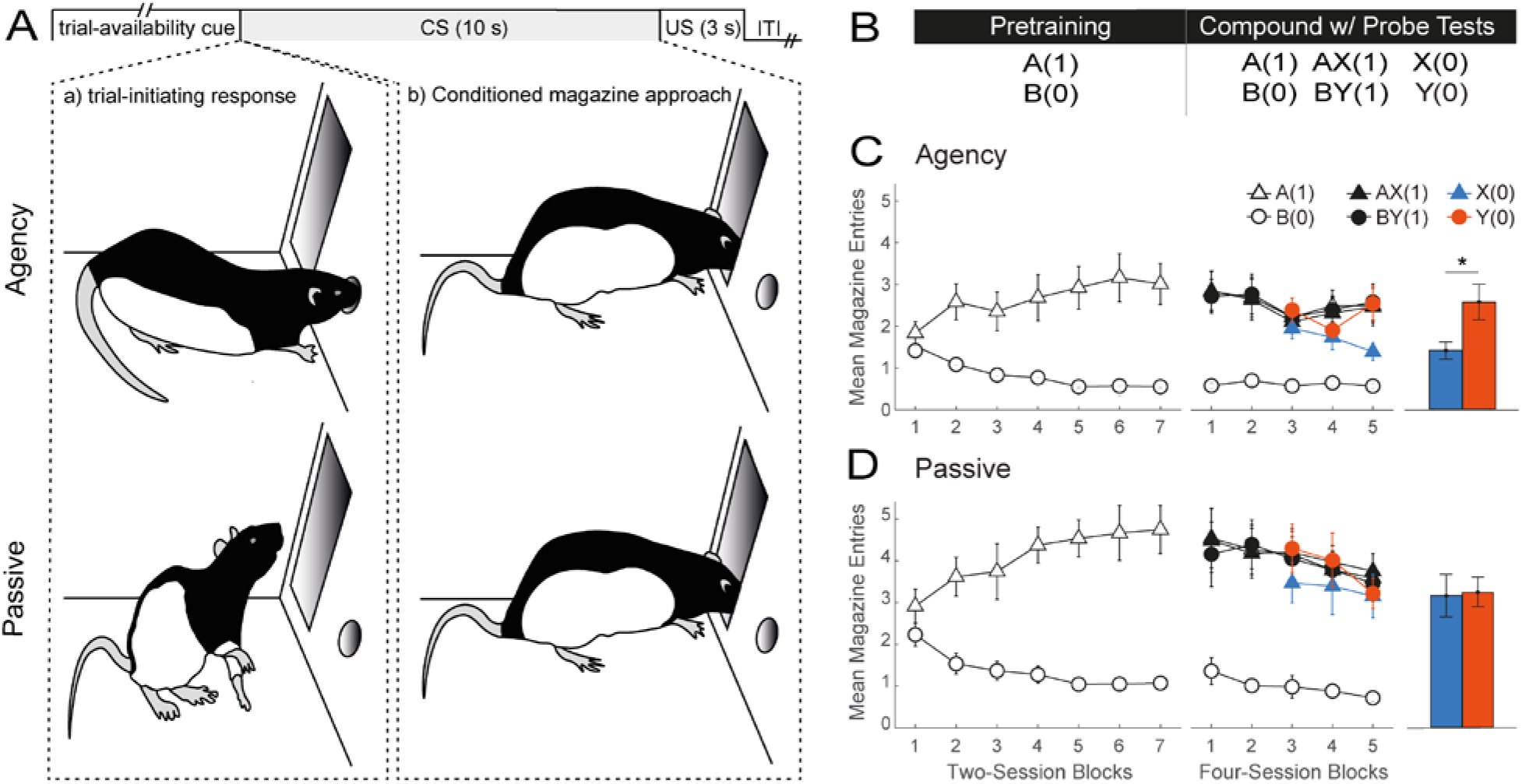
Agency over learning rescues competitive credit assignment from the deleterious effects of massed training in a blocking task. **(A)** Trial structure in this and the remainder of the studies. On each trial offer, a nose port light was presented in both groups signaling trial availability to Agency rats. A trial-initiating response (nose poke) by an Agency rat immediately resulted in a 10-s cue being presented to that rat as well as to its yoked animal in the Passive group. On reinforced trials, a sucrose US was presented at dipper magazine following cue offset. Trial offers were separated by a 10-s variable intertrial interval (ITI). **(B)** Experimental design. Letters A and B denote visual stimuli, whereas X and Y denote auditory stimuli. Digits in brackets represent the probability of reward for each trial type. The pretraining phase involved a simple discrimination between A and B. During the compound phase, these trials were interleaved with compounds AX (where A should block X) and BY (where B should not block Y), both continuously reinforced. To test for blocking (i.e., less responding to X than Y), two daily probe trials with X and Y were introduced on session 9 of the Compound phase. **(C)** and **(D)** Behavioral results in groups Agency and Passive, respectively. The left and center line plots depict performance during the Pretraining and Compound phases, respectively. The bar graphs on the right show average responding to X and Yon probe trials across the last four sessions. Conditioned responding is measured as mean number of head entries (+/− SEM).

In both groups, a sucrose reward delivered in a dipper cup was made available on reinforced trials immediately after the termination of the cues. Conditioned responding was measured as the number of anticipatory head entries made by the rat at the dipper recess during the last 5 s of cue presentation (Holland 1977; see Materials & Methods). Critically, the ITI was programmed to be only 10 s on average (range: 5-15 s). Since Agency rats could forgo trial offers, the mean ITI was effectively longer (see Materials & Methods), but still considerably shorter than the mean ITI typically used in studies of conditioned magazine approach featuring 10-s cues (in the order of minutes).

Both groups underwent two phases of training. In the first phase, rats were pretrained with a simple discrimination involving two visual cues, A and B. Cue A, which would serve as the blocking stimulus in the following phase, was reinforced with a probability of 1 [henceforward symbolized by A(1)], while B was never reinforced [B(0)]. Training continued for 14 days to allow the opportunity for asymptotic discrimination learning (Fig. 1, Panels C and D, left). A mixed ANOVA revealed a significant main effect of cue (F_(1,182)_ = 398.24, p < 0.001) and group (F_(1,14)_ = 8.10, p = 0.013), and a group by cue interaction (*F*_(1,182)_ = 15.65, *p* < 0.001). Post-hoc analyses revealed that this interaction was likely driven by the slightly higher level of responding on A(1) trials in the Passive group (*t*_(17.1)_ = −3.95, *p* = 0.006), as both groups significantly discriminated between A(1) and B(0) (Agency: *t*_(182)_ = 11.31, *p* < 0.001; Passive: *t*_(182)_ = 16.91, *p* < 0.001).

In the second, compound phase (Fig. 1, Panels C and D, center), training with A(1), B(0) continued for another 20 sessions, but, in addition, two novel auditory cues, X and Y, were presented in compound with A and B on separate, reinforced trials. Specifically, X accompanied A as the stimulus to be blocked, whereas Y accompanied B as the control cue for blocking [AX(1), BY(1)]. Panels C and D (center) of Fig. 1 show that in both groups the compounds evoked similar levels of conditioned responding as A. A mixed ANOVA revealed a main effect of cue (*F*_(3,266)_ = 187.35, *p* < 0.001), session block (*F*_(4,266)_ = 4.28, *p* = 0.0002) and group (*F*_(1,14)_ = 7.07, *p* < 0.019), and a group by cue interaction (*F*_(3,266)_ = 10.33, *p* < 0.001). Post hoc analyses revealed no significant between-group differences for any of the trial types, indicating that, once again, the interaction was likely driven by the higher asymptote of responding in the Passive group on reinforced trials.

In order to monitor the emergence of blocking (i.e., less responding to X than Y), two probe trials with each of X and Y were randomly interleaved daily from session 9 onward (Fig. 1, panels C and D, center). Inspection of the results suggests that a blocking effect emerged at the end of the compound phase in the Agency, but not the Passive group. This impression was confirmed by a mixed ANOVA that focused on the mean responding to X and Y across the last four probe sessions (Fig. 1, panels C and D, right). This analysis revealed significant main effects of cue (*F*_(1, 98)_) = 7.80, *p* = 0.006) and group (*F*_(1, 14)_) = 6.42, *p* = 0.024), and a significant group by cue interaction (*F*_(1,98)_ = 6.28, *p* = 0.014). Exploration of this interaction with simple main effects confirmed a significant difference in responding to the cues (i.e., an intermixed blocking effect) in the Agency (*t*_(98)_ = 3.75, *p* = 0.002), but not the Passive group (*t*_(98)_ = 0.20, *p* = ∼1). The results thus provide evidence that agency over learning rescues competitive credit assignment from the adverse effects of massed trials.

### Agency rescues competitive credit assignment in a novel cue competition task

To further examine the influence of agency on competitive credit assignment under massed trials, we next compared the performance of Agency and Passive groups in a novel cue competition design. This design creates a conflict between the expected pattern of responding to two cues, X and Y, when credit assignment is competitive relative to when it is noncompetitive. By creating this conflict, this design maximizes the chances of detecting differences between competitive and noncompetitive learning. This makes this design ideally suited for examining the full impact of behavioral and neural manipulations on credit assignment. Given the novelty of the design, we began by piloting it in a standard Pavlovian magazine-approach setting with spaced trials (Supplementary Materials, Exp. S1).

The details of the experimental design are shown in the table of Fig. 2A. Two groups were trained with the same master-yoked procedure used in the previous study (Fig. 1A). In the pretraining phase (Fig. 2, panels B and C, left), rats received 10 sessions of discrimination training with two visual cues, A(1) and B(0), and two auditory cues, X(.75) and Y(.25) where, once again, the numbers in parenthesis represent the probability of reward associated with each cue. A mixed ANOVA revealed a main effect of cue (*F*_(3,266)_ = 24.27, *p* < 0.001) and a cue by session block interaction (*F*_(12,266)_ = 2.03, *p* = 0.022). Post-hoc analysis of this interaction revealed that the discrimination between A(1) and B(0) was solved from session block 3 onward (*t*_(266)_ = [3.78 – 4.64], *p* < 0.037). Post hoc analysis of the main effect of cue revealed that, overall, responding to X(.75) was higher than to Y(.25) (*t*_(266)_ = 5.25, *p* < 0.001). No effect of group nor any interaction involving that factor was found.

**Figure 2.**
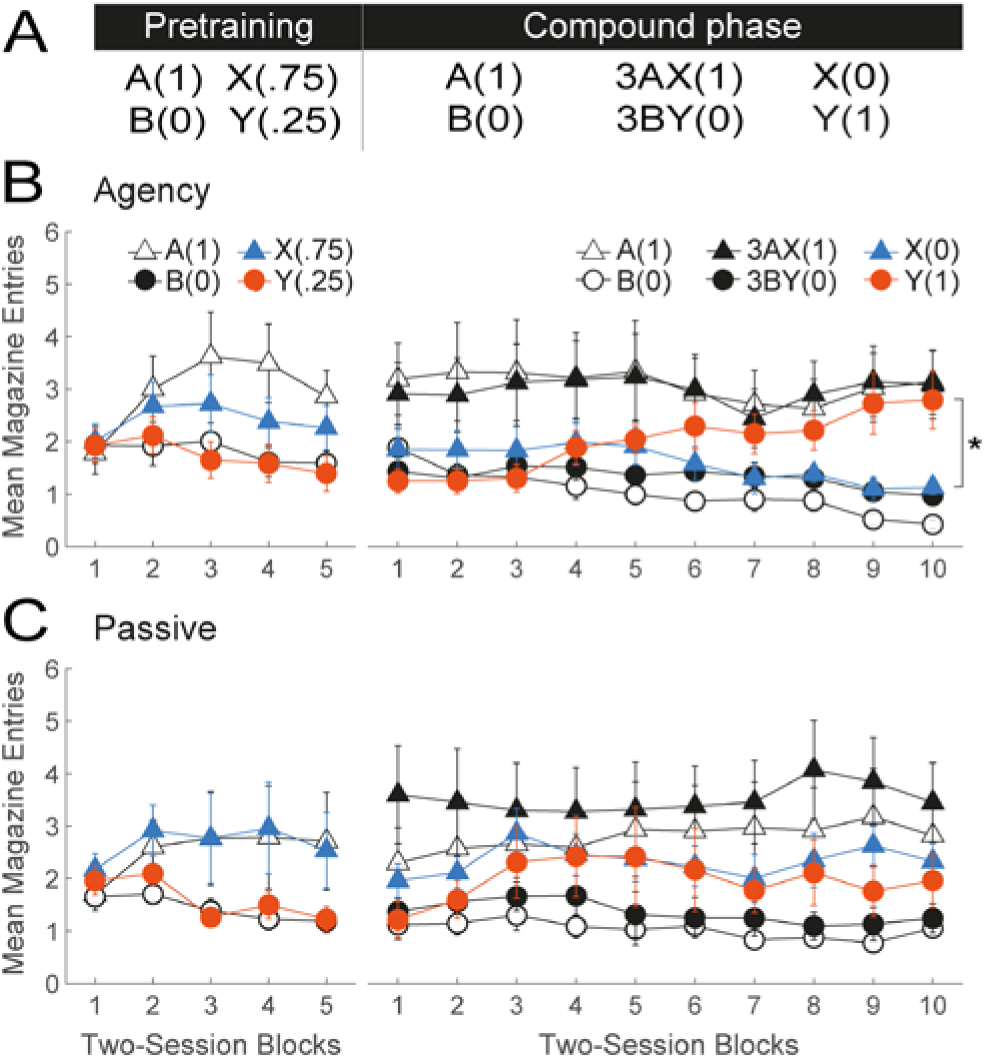
Agency over learning rescues competitive credit assignment from the deleterious effects of massed training in a novel cue-competition task. **(A)** Experimental design. The coefficient 3 denotes three times as many presentations of the AX(1) and BY(O) compounds as of the rest of trial types. During the Pre-training phase, all rats received discrimination training between A and B and X and Y. In the Compound phase, these stimuli continued to be presented with the same probability of reward. However, X was presented in compound with A on the 75% of trials in which it was rewarded (allowing A to take away its credit), whereas Y was presented in compound with B on the 75% of trials in which it was not rewarded (allowing B to take the blame for reward omission). Trials with X(O) and Y(1) permitted continual monitoring of the predictive status of these stimuli as the discrimination developed. **(B)** and **(C)** Behavioral results in groups Agency and Passive, respectively. The left and right line plots depict performance during the Pretraining and Compound phases, respectively. Conditioned responding is represented as mean number of magazine head entries (+/− SEM).

In the second, compound phase, cues X and Y had the same probability of reward, but were subject to opposing competing forces (Fig. 2, Panels B and C, right). Specifically, X was presented in compound with A on the 75% of trials in which it was followed by reward [3AX(1), X(0); where 3 indicates the proportion of trials]. This allowed A to compete with X as a predictor of reward and *steal* its credit (e.g., Wagner, 1969). A’s ability to serve as a competitor was further bolstered by continuing to present it by itself followed by reward [A(1)]. In addition, cue Y was presented in compound with B on the 25% of trials in which Y was not reinforced [3BY(0), Y(1)], allowing B to compete with Y for predicting reward omission. Casually put, this training was intended to ensure that B rather than Y would *take the blame* for the omission of reward on 3BY(0) trials (Chorazyna, 1962; Rescorla, 2003). Throughout this phase, B(0) trials continued to be presented.

A key advantage of this design is that X(0) and Y(1) trials permit online monitoring of the impact of competition on responding to these cues. If credit assignment is noncompetitive (e.g., Bush & Mosteller, 1951), X should be expected to evoke more responding than Y given its higher probability of reward. Conversely, to the extent credit assignment is competitive, Y should be expected to evoke more responding than X (Wagner et al., 1968; Wagner, 1969). To examine the role of agency, we focused our analysis on responding on X(0) and Y(1) trials. Inspection of Fig. 2 (Panels B & C, right) suggests that cue competition prevailed in the Agency, but not the Passive group. This impression was confirmed by a mixed ANOVA, which revealed significant group by cue (*F*_(1, 266)_ = 14.69, *p* < 0.001), cue by session block (*F*_(9, 266)_ = 2.19, *p* < 0.001), and group by cue by session block interactions (*F*_(9, 266)_=2.16, *p*=0.025). A post-hoc analysis of the three-way interaction revealed that, consistent with competitive credit assignment, rats in the Agency group responded to Y significantly more than to X on session blocks 9 (*t*_(266)_ = 3.69, *p* < 0.002) and 10 (*t*_(266)_=3.78, *p* < 0.002). In contrast, in the Passive group, the difference between X and Y was marginally significant only on session block 9, but in the opposite, noncompetitive direction (X > Y) (*t*_(266)_ = 1.97, *p* = 0.05).

A mixed ANOVA on responding to the remainder of the cues in the compound phase revealed a significant main effect of cue (*F*_(3,546)_ = 138.82, *p* < 0.001) and a group by cue interaction (*F*_(3,546)_ = 3.23, *p* = 0.022). Post-hoc analyses, however, confirmed that that both groups discriminated between A(1) and B(0) [Agency: *t*_(546)_ = 10.81, *p* < 0.001; Passive: *t*_(546)_ = 9.26, *p* < 0.001] as well as between 3AX(1) and 3BY(0) trials [Agency: *t*_(546)_ = 8.79, *p* < 0.001; Passive: *t*_(546)_ = 11.42, *p* < 0.001]. A likely contributor to this interaction was the greater responding observed on 3AX(1) than A(1) trials in the Passive (*t*_(546)_ = −3.86, *p* = 0.004), but not the Agency (*t*_(546)_ = 0.47, *p* = 1) group. This difference can be explained by the fact that X undergoes limited competition by A in the Passive group, allowing the two cues to sum their predictive credits when presented in compound. This study thus provides further evidence of a profound impairment in cue competition under massed Pavlovian training, rendering the rescuing effects of agency over learning the more striking. One interpretation, however, is that Agency rats did not apportion credit any more competitively than Passive rats, but instead treated cues X and Y as radically distinct when presented alone vs. in compound. In other words, Agency, but not Passive rats may have treated the compounds AX and BY as independent *configural* stimuli distinct from their constituent elements. To rule out this interpretation, we used a variant of the above design that does not afford an explicit configural solution.

### Agency rescues competitive credit assignment in the absence of an explicit configural solution

Fig. 3A shows the experimental design. In the pretraining phase, all rats received 8 sessions with a simple visual discrimination of the form A(1), B(0) (Fig. 3, panels B and C, left). Both groups solved this discrimination as evidenced by greater magazine-approach responding on A(1) than B(0) trials over the course of training (main effect of cue: *F*_(1,98)_ = 88.72, *p* < 0.001; cue by session block interaction: (*F*_(1,98)_ = 5.32, *p* < 0.002). No significant effects of group or interactions involving that factor were found.

**Figure 3.**
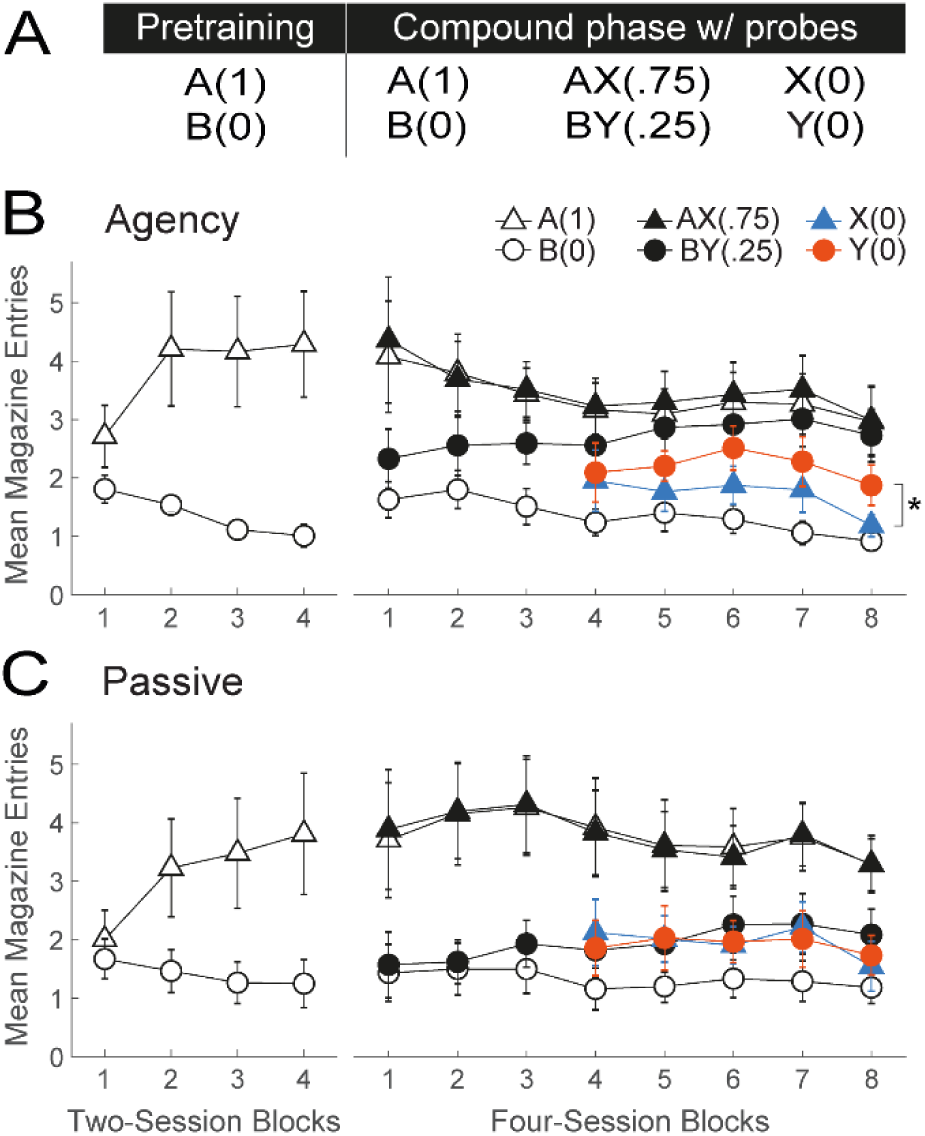
Agency over learning rescues competitive credit assignment under massed training conditions in the absence of an explicit configural solution. **(A)** Experimental design. The pretraining phase involved a simple discrimination between A(1) and 8(0). During the compound phase, these trials continued to be presented, but were interleaved with AX(.75) and BY(.25) trials. Note that X has a higher probability of reward than Y. However, X signals a net decrement in the probability of reward when considered against the backdrop of A, whereas Y signals a net increase in the probability of reward when considered against the backdrop of Y. Thus, to the extent credit assignment is competitive, Y should evoke more responding than X, but the opposite should be true if credit assignment is noncompetitive. To test this, two daily probe trials with X and Y were inter-leaved with training trials, starting on session 13 of the Compound phase. **(B)** and **(C)** Behavioral results in groups Agency and Passive, respectively. The left and right line plots depict performance during the Pretraining and Compound phases, respectively. Conditioned responding is represented as mean number of magazine head entries (+/− SEM).

In the second, compound phase, all rats continued to receive the A(1), B(0) discrimination, but novel compound trials AX(.75) and BY(.25) were introduced, where X and Y were again auditory stimuli (Fig. 3, panels B and C, right). A mixed ANOVA on responding to A(1), B(0), AX(.75) and BY(.25) trials throughout this phase revealed a significant effect of cue [*F*_(3,434)_ = 142.33, *p* < 0.001] and a group by cue interaction [*F*_(3,434)_ = 7.88, *p* < 0.001]. Post-hoc analyses of this interaction revealed that both groups solved the A(1) vs. B(0) [Agency: *t*_(434)_ = 10.94, *p* < 0.001; Passive: *t*_(434)_ = 13.28, *p* < 0.001] as well as the AX(.75) vs. BY(.25) [Agency: *t*_(434)_ = 4.36, *p* < 0.001; Passive: *t*_(434)_ = 9.96, *p* < 0.001] discriminations. A likely contributor to the group by this interaction is the fact that Agency rats discriminated better between BY(.25) and B(0) trials [*t*_(434)_ = −7.21, *p* < 0.001, Cohen’s d = 0.79] than Passive rats [*t*_(434)_ = −3.30, *p* = 0.029, Cohen’s d = 0.53]. This suggests that cue B was better able to protect Y from extinction in Agency but not Passive rats, consistent with the notion that competition was impaired in the latter.

Note that, in the absence of competitive credit assignment, this training should result in X evoking more responding than Y given their respective probabilities of reward (0.75 and 0.25). On the other hand, if credit assignment is competitive, more responding to Y than X should be observed. This is because nonreinforced AX trials should turn X into a signal for the occasional omission of reward (a conditioned inhibitor), while the presence of B on nonreinforced BY trials should protect Y from extinction. To test this, we randomly interspersed two daily nonreinforced probe trials with X and Y [X(0), Y(0)] starting on session 13 (Fig. 3, panels B and C, right). A mixed ANOVA on responding during the probe trials revealed a marginally significant effect of cue (*F*_(1,126)_ = 3.86, *p* < 0.052) and a significant group by cue interaction (*F*_(1,126)_ = 5.43, *p* < 0.021). Post-hoc analysis of this interaction confirmed that the Agency group responded significantly more to Y than X (*t*_(126)_ = −3.04, *p* = 0.017). In contrast, the Passive group responded equally to both cues (*t*_(126)_ = 0.26, *p* = 1), as expected if cue competition was disrupted. The results thus provide further evidence that agency over learning rescues competitive credit assignment.

### Ruling out alternatives for the role of agency in competitive credit assignment

The results so far can be readily interpreted by assuming that agency over learning enhances the animals’ attention to task, discriminating proficiency, or ability to process compounded stimuli concurrently. To test these interpretations, we compared performance between Agency and Passive rats in a patterning task in which opposite credit must be assigned to compound cues and their constituent elements (Fig. 4A). One such problem was a negative-patterning discrimination in which two cues, a visual (A) and an auditory (X) stimulus, were rewarded when presented individually, but not in compound [A(1), X(1), AX(0)]. A second problem involved a positive-patterning discrimination in which another pair of visual (B) and auditory stimuli (Y) were rewarded when presented in compound, but not individually [B(0), Y(0), BY(1)]. If any of the aforementioned interpretations is correct, Passive animals should find this discrimination particularly difficult.

**Figure 4.**
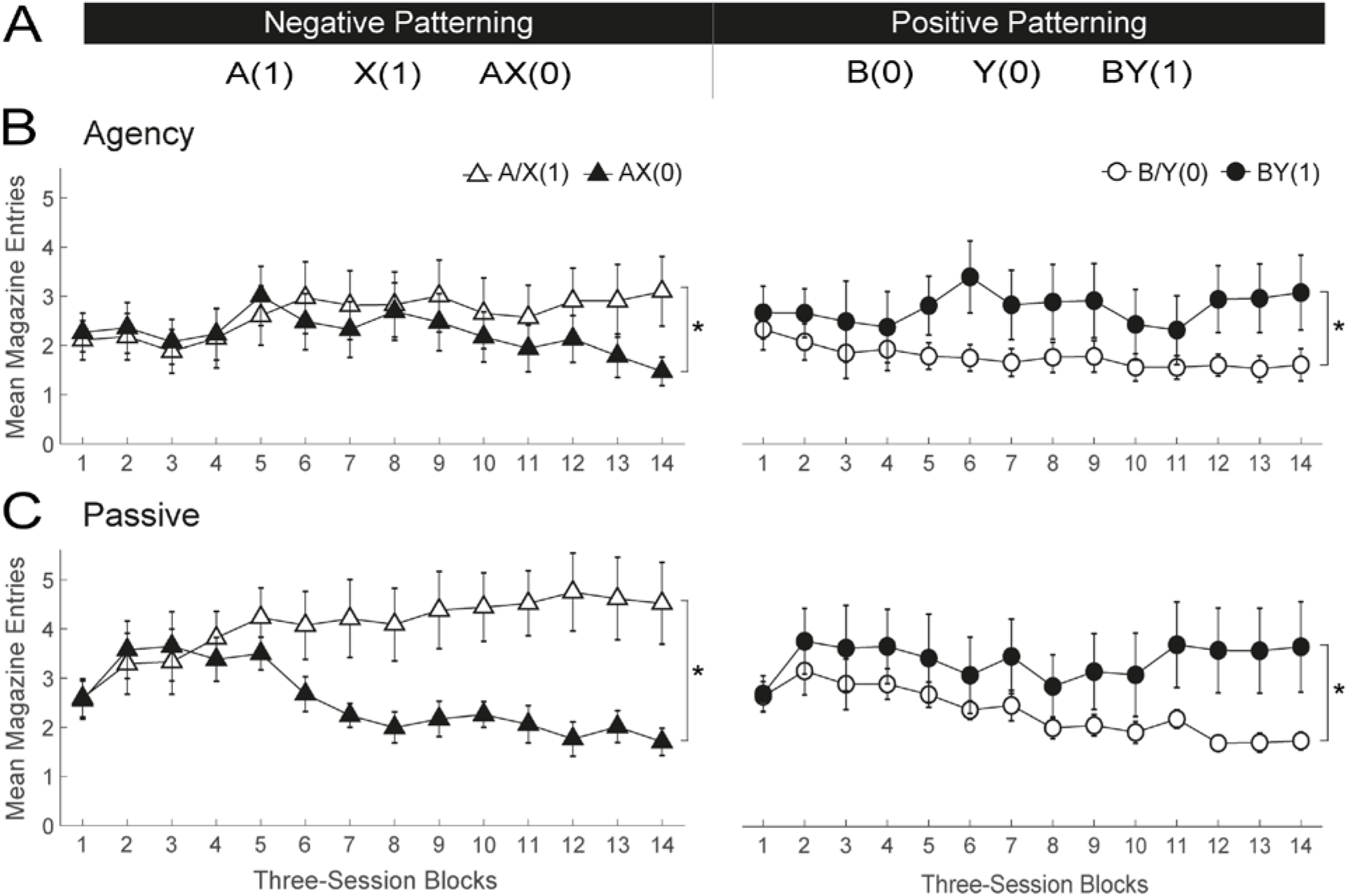
Passive rats perform no worse than Agency rats in a patterning task, suggesting spared compound processing and general discrimination ability. **(A)** Experimental design. A negative- and positive-patterning discrimination involving two visual stimuli (A and B) and two auditory stimuli (X and Y) were trained contemporaneously. **(B)** and **(C)** Behavioral results in groups Agency and Passive, respectively. The left and right panels show discrimination performance in the negative- and positive-patterning discriminations, respectively. Note that, for each discrimination, responding has been averaged across elemental trials [e.g., A/X(1) denotes mean responding on A(1) and X(1) trials]. Conditioned responding is measured as the mean number of magazine head entries (+/− SEM).

Although both discriminations were trained concurrently, for simplicity’s sake we treated them separately when displaying and analyzing the data (Fig. 4, Panels B and C). To further simplify the analysis, we averaged responding on elemental trials [mean of A(1) and X(1) trials and of B(0) and Y(0) trials]. Inspection of the data suggests that Passive animals solved both patterning problems at least as well as Agency rats. A mixed ANOVA on the negative-patterning discrimination revealed a main effect of group [*F*_(1,378)_ = 91.60, *p* < 0.001] and significant session block by cue [*F*_(13,378)_ = 5.11, *p* < 0.001] and group by cue [*F*_(1,378)_ = 33.48, *p* < 0.001] interactions. Simple main effects analysis of the latter interaction confirmed that both Agency [*t*_(378)_ = −2.68, *p* = 0.016] and Passive [*t*_(378)_ = - 10.86, *p* < 0.002] animals solved the A(1)/X(1) vs. AX(0) discrimination, although the effect size was larger in Passive than Agency rats (Cohen’s d = 0.93 and 0.23, respectively). A parallel mixed ANOVA of the positive-patterning discrimination revealed only a significant effect of cue [*F*_(1,378)_ = 106.56, *p* < 0.001], indicating that both groups solved the B(0)/Y(0) vs. BY(1) discriminations similarly (Agency: Cohen’s d = 0.65; Passive: Cohen’s d = 0.68). Taken together, these findings suggest that the deficits in competitive credit assignment previously observed in Passive rats were unlikely due to an inferior level of engagement, ability to solve complex discriminations, process compounded stimuli concurrently, or form configural representations.

To buttress these conclusions, we also presented this patterning problem to the rats from the first study at the end of blocking training (Supplementary Materials, Exp. S2). Since those rats had already experienced the two auditory stimuli as X and Y, two novel auditory stimuli were used. The results of this replication confirmed those with naïve animals. That is, the same Passive rats that exhibited deficits in blocking were at least as capable of solving complex nonlinear discriminations as their Agency counterparts.

## General Discussion

Competitive credit assignment is the backbone of models of associative and reinforcement learning, to the point that an inability to account for competitive learning phenomena renders a model obsolete and useless (e.g. Ludvig et al., 2012; Miller et al., 1995). Yet converging evidence indicates that competition is not automatically determined by the presence of other cues, but also by learning conditions such as trial spacing (reviewed in: Urcelay, 2017; Witnauer et al., 2014; Wheeler & Miller, 2008). Specifically, when information is presented in a massed fashion, cue competition is diminished (Stout et al., 2003; Sissons & Miller, 2009; Packheiser et al., 2020; Beesley & Shanks, 2012). Here, we provide evidence that the presence of agency over the learning experience can rescue competitive credit assignment from the detrimental effects of massed training. This evidence speaks to the necessity of incorporating agency into current theories of learning. Taken together, our data allow us to rule out various trivial explanations for the effects observed. Firstly, the beneficial effects of agency on competitive credit assignment are not simply the product of a heightened ability to process compounded stimuli concurrently or learn complex discriminations. Evidence for this comes from the superior performance of Passive groups in the patterning task. Secondly, neither can the contribution of background (contextual) cues to competitive learning explain our full pattern of results. While contextual conditioning could summate with responding to the target cues (Urcelay & Miller, 2014) and mask any differences when responding is at ceiling (e.g., Passive group in blocking study), such masking would not occur when responding is below asymptote (e.g., in novel cue-competition task studies). Thirdly, our data are likewise difficult to explain by a differential role of eligibility traces in Agency and Passive groups (Sutton, 1988; Sutton & Barto, 2018). Specifically, in Passive animals, massed training might allow eligibility traces of recently presented cues to spill over the subsequent trial and contaminate credit assignment. This effect would be weaker in Agency rats if trial-initiating responses serve to precipitate the decay of eligibility traces, essentially fulfilling the role of a long ITI. The issue with this hypothesis is that it also predicts poorer performance for Passive rats in the patterning task, which our data disconfirmed.

The effects observed can be explained by assuming that agency over learning modulates the computation of prediction errors (PEs; i.e., the difference between actual and expected outcome). The standard assumption in models of associative and reinforcement learning is that PEs are computed by summing across the outcome expectancies generated by each of the cues present (Rescorla & Wagner, 1972; Sutton & Barto, 1990; Siegel & Allan, 1996; Sutton & Barto, 2018; Honey et al., 2020). This aggregate expectancy is then compared with the actual outcome experienced, and the difference determines the size and sign of the credit update (i.e., learning). An aggregate PE implies that all cues present share a common engine of learning which ensures that they compete over a common pool of available credit. Our data suggest that, in the absence of agency, massed training might interfere with the computation of aggregate PEs and render learning less competitive. This might explain why the patterning task was solved more readily by Passive rats, as cue competition can hinder the solution of nonlinear discriminations (e.g., Byrom & Murphy, 2018). Given the central role of PEs in learning (e.g., Schmajuk, 2010; Sutton & Barto, 2018), cognition (e.g., de Bruin & Michael, 2017; den Ouden & de Lange, 2012), psychopathology (e.g., Krawczyk et al., 2017; Zald & Treadway, 2017; Gradin et al., 2011) and neuroscience (e.g., Diederen & Fletcher, 2020; Watabe-Uchida et al., 2017; Ohmae & Medina, 2015; Roesch et al., 2012; Niv & Schoenbaum, 2008; Waelti et al., 2001), further examination of its interaction with agency would be of fundamental importance.

Importantly, agency over learning might regulate other mechanism besides PE; for instance, by modulating the allocation of attention to cues (e.g., Mackintosh, 1975; Pearce & Hall, 1980; Krushke, 2001; Le Pelley, 2004; Esber & Haselgrove, 2011; George & Pearce, 2012). According to a classical selective attention account (Mackintosh, 1975), cue competition results from paying increasingly more attention to the best predictor while devoting increasingly less attention to relatively poor predictors. This attentional divergence ensures that the most reliable cue acquires substantial credit and controls behavior while limiting the ability of unreliable cues to do so. Agency over learning might thus protect the attentional mechanism underlying the divergence from detrimental effects of massed training. Alternatively, the effect of agency on credit assignment need not be limited to learning, but, rather, could work at the level of memory retrieval through a comparator mechanism (Denniston et al., 2001; Stout & Miller, 2007).

The current findings have far-reaching implications both for normal functioning and mental health. In pedagogical settings, where massed instruction has been long shown to be detrimental (Ebbinghaus, 1885; Grote, 1995; Seabrook et al., 2005; Rohrer, & Taylor, 2006; Moulton et al., 2006; Patocka et al., 2019; Namaziandost et al., 2020), our findings raise the possibility that agency over the presentation of information might promote more selective, discriminating encoding and retention. In the context of mental health, it might open opportunities for therapeutical interventions based on enhancing the individual’s perceived sense of agency in disorders characterized by attentuated selective learning, including psychotic (Jones et al., 1992; Moran et al., 2003; Fletcher & Frith, 2009), attentional (Oades & Müller, 1997), anxiety (Boddez et al., 2012), and substance use disorders (SUDs) (Freeman et al., 2013; Muscat et al., 2008; Muscat & Spiteri, 2011).

For the present, much work is needed to elucidate the complex role that agency is likely to play in learning and psychopathology. Consider the case of SUDs, where agency is known to mitigate some of the more dramatic and aversive effects of drugs of abuse (Twining et al., 2009; Weise-Kelly & Siegel, 2001; Dworkin et al., 1995). This role might in part result from credit assignment being more competitive, which should prevent incidental and redundant stimuli from contributing to cue reactivity. Our results suggest that as the sense of agency over drug consumption wanes and drug-related behaviors transition from voluntary and goal-directed to habitual and compulsive (Everitt & Robbins, 2005; 2016), credit assignment might also become less competitive. This transition would exacerbate the individual’s vulnerability to drug abuse and relapse by drastically expanding the set of stimuli capable of inducing cue reactivity. Tantalizingly, long-term exposure to potent rewards such as cocaine, heroin, and sucrose has been shown to impair competitive credit assignment (Lucantonio et al., 2015; Sharpe et al., 2015). In light of such implications, the present findings call for a closer investigation of the role of agency in credit assignment.

## Materials and Methods

For the sake of convenience, the four studies above will be referred to in this section as Exps. 1-4, and correspond, respectively, to the blocking task (Fig.1), the novel cue-competition task (Fig. 2), its second variant (Fig. 3), and the patterning task (Fig. 4).

### Subjects

All studies used 16 experimentally-naïve, gender-balanced, adult Long-Evans rats, making a total of 64 animals. The age and weights of the rats at the outset of each experiment was as follows. In Exp. 1, rats were ∼20 weeks old (wo) and weighed 441-516 g (males) and 257-290 g (females); in Exp. 2, rats were ∼13 wo and weighed 342-388 g (males) and 234-269 g (females); in Exp. 3, rats were ∼20 wo and weighed 448-529 g (males) and 269-298 g (females); in Exp. 4, rats were ∼22 wo and weighed 475-554 g (males) and 284-325 g (females). All animals were bred at Brooklyn College from commercially available populations (Charles River Laboratories). They were housed individually in standard clear-plastic tubs (10.5 in. × 19 in. × 8 in, Charles River Laboratories) with woodchip bedding. The colony room was maintained on a 14:10 light/dark cycle schedule. Behavioral sessions were conducted between 7-10 hours after the onset of the light phase of the cycle. Throughout training, water access was restricted to 1 h/day following each experimental session while food was provided *ad libitum*. All animal care and experimental procedures were conducted according to the National Institutes of Health’s *Guide for the Care and the Use of Laboratory Animals*, and approved by the Brooklyn College Institutional Animal Care and Use Committee (Protocol #303).

### Apparatus

Behavioral training was conducted in eight modular conditioning chambers (32-cm long X 25-cm wide X 33-cm tall, Med Associates, Inc.). Each chamber was enclosed in a ventilated sound-attenuating cubicle (74 cm x 45 cm x 60 cm) fitted with an exhaust fan that provided a background noise level of ∼50 dB. All reported locations of stimulus and response apparatus were measured from the grid floor of the conditioning chamber to the lowest point or edge of the apparatus. The left wall of the chamber housed two white jewel lamps (28V DC, 100 mA) mounted on the left and right panels 9.3 cm from the grid floor. Above each of these lamps was a speaker located 20.6 cm above the grid floor and connected to a dedicated tone generator capable of delivering a 2.5-Hz, 80-dB clicker (left panel) and a 70-dB white noise (right panel). Two additional speakers were located on the left and right panels of the right wall of the chamber 24.8 cm above the grid floor. Each of them was also connected to a dedicated speaker capable of delivering a 12-kHz, 70-dB tone (left panel) and a 1-kHz, 80-dB tone (right panel). The right wall also housed a third jewel lamp located on the center panel 17.2 cm above the grid floor. Below this lamp, 4.6 cm above the grid floor, was a circular noseport 2.6 cm in diameter, equipped with a yellow LED light and an infrared sensor for detecting nose entries. This noseport was flanked by a recessed liquid reward magazine (aperture: 5.1 cm x 15.2 cm) located on the right panel, 1.6 cm above the grid floor. This magazine was equipped with an infrared sensor for detecting head entries, and connected to a liquid dipper that could deliver a 0.04 cc droplet of a 10% sucrose solution. The chambers remained dark throughout the experimental session except during presentations of the visual stimuli. In the same room was a computer running Med PC IV software (Med Associates Inc., St. Albans, VT, USA) on Windows OS which controlled and automatically recorded all experimental events via a Fader Control Interface.

### Procedure

#### Magazine training

Prior to the beginning of each study, rats were first randomly assigned to either the master or yoked group—labeled groups Agency and Passive, respectively—with the constraint that each group be gender-balanced. Each animal assigned to the Agency group was paired with an age and sex-matched Passive group animal. All sessions began with a 2-min acclimation period in the conditioning chambers. Rats initially received a session of magazine training in which they learned to retrieve a sucrose reward from the dipper cup. This session lasted 62 min and consisted of 60 trials. For the first 10 trials, sucrose was made available for 30 s every 30 s; for the second 20 trials, it was available for 20 s every 40 s; and for the last 30 trials, it was available for 10 s every 50 s.

#### Shaping

In all four studies, Agency rats went on to receive five shaping sessions in which they learned to self-initiate trials, following the procedure developed by Reverte at al. (2020). On the first shaping session, the noseport light was turned on for a maximum of 20 s, during which a nose poke at the nose port immediately resulted in the termination of the noseport light and a 10-s period of sucrose availability. Trials were separated by a 10 s variable ITI (range: 5−15 s). Failure to respond at the nose port resulted in the noseport light coming off and the trial being repeated after a regular ITI. Over the following four shaping sessions, we introduced and progressively increased a delay of 2, 4, 6, and 8 s between the rat’s response at the port and sucrose availability. During this delay, the noseport light would flash at a 1-Hz frequency (on for 0.5 s, off for 0.5 s). Concurrently, reward availability was progressively shortened (8, 6, 4, and 3 s). Throughout shaping training (and for the remainder of the experiment), Passive were yoked to their Agency counterparts to ensure that they received the same exact sequence of events and at the same time, except, of course, for the trial-initiation response.

#### Trial structure

The trial structure was common to all four studies (Panel A, Fig. 1). Following shaping, experimental sessions began with a 30-s acclimation period. Agency rats would then be presented with their first opportunity to start a trial as signaled by trial-availability cue (the onset of the noseport light). The duration of this cue was 20 s, during which a response at the noseport would immediately turn off the noseport light and turn on one of various possible visual, auditory, or audiovisual compound cues which were always 10 s long. The trial types specified by these cues were selected from a pseudorandom list built with the constraint that no trial type could be presented more than three times in succession. On reinforced trials, the cues’ offset coincided with the delivery of a 0.04-cc bolus of sucrose, which remained available for 3 s, after which a short ITI followed (mean: 10 s; range: 5-15 s). As during shaping, failure to self-initiate a trial terminated the noseport light after 20 s and led to a regular ITI period (mean: 10 s; range: 5-15 s). Passive rats received the same sequence of events—including the same trial types at the same time and in the same order—as their Agency counterparts, but in standard Pavlovian fashion (i.e., noncontingent on any response). For any yoked pair of rats, a session terminated once the Agency rat completed all scheduled trials or timed out after 90 min.

Since Agency rats self-paced their training, they could take breaks that elongated the effective average ITI. Such pauses were of course also applied to their yoked counterparts. Supplementary Table S1 provides the effective ITI durations as well as the total session durations for each experiment. Furthermore, granting agency over trial presentation necessarily entailed the risk that rats would not complete all scheduled trials within the imposed time limit of 90 min, in which case the session would time out. Once again, the use of a master-yoked procedure ensured that this issue affected both groups equally. Supplementary Table S2 lists all incomplete sessions for all studies presented.

#### Discrimination training

##### Experiment 1

Training comprised two phases (see table in Panel B, Fig. 1). In the first, pretraining phase, rats in both groups received 14 sessions of A(1) vs. B(0) discrimination training, where A and B were visual cues and the numbers in parenthesis represent the probability of reward. One visual cue was constructed by flashing the two jewel lamps on the left wall alternately at a 2-Hz frequency (on for 0.25 s, off for 0.25 s), whereas the other was provided by the steady illumination of the white jewel lamp located on the right wall. These cues were counterbalanced, and were presented 48 times each in a session.

The second, compound phase comprised 20 sessions, during which rats continued to receive A(1), B(0) trials presented 36 times each per session. In addition, compound trials AX(1) and BY(1) trials were introduced, where X and Y represent two auditory cues. These auditory cues were provided by a 12-kHz, 70-dB tone and a 70-dB white noise, counterbalanced. There were 12 presentations of each of the AX and BY compounds per session. From session 9 to the end of the compound phase, two probe trials with each of the target cues, X (i.e., the cue to be blocked) and Y (the control cue) were additionally administered. This increased the total number of trials in Phase 2 from 96 to 100.

##### Experiment 2

Training consisted of two phases (see table in Panel A, Fig. 2). In the first, pretraining phase, rats received 10 sessions of A(1) vs. B(0) and X(.75) vs. Y(.25) discrimination training, where A and B were the same visual cues and X and Y were the same auditory cues used in Exp. 1, also counterbalanced. Once again, the numbers in parenthesis indicate the probability of reward. Each cue was presented 24 times in a session.

In the second, compound phase, the probability of reward for each cue trained in Phase 1 was maintained constant, but cue A was added to all trials in which X was reinforced, whereas B was added to all trials in which Y was not reinforced. Thus, the compound phase consisted of the following trial types: 10A(1), 10B(0), 30AX(1), 10X(0), 30BY(0), 10Y(1), where the coefficients represent the number of trials presented in a session (100 trials in total). Phase-2 training proceeded for 20 sessions.

##### Experiment 3

The study comprised two phases (see table in Panel A, Fig. 3). In the pretraining phase, rats received 8 sessions of A(1) vs. B(0) discrimination training, where A and B were the same visual cues used in Exp. 1. Each of these trial types was presented 48 times in a session.

In the second, compound phase, rats received 32 sessions of discrimination training. During this phase, A(1) vs. B(0) training continued, but audiovisual compounds AX(.75) and BY(.25) were added, with X and Y being the same auditory stimuli used in Exp. 1. Specifically, the compound phase consisted of the following training trials: 24A(1), 24B(0), 18AX(1), 6AX(0), 6BY(1), 18BY(0), where the coefficients denote the number of trials presented in a session. Starting on session 13, two probe trials with cues X and Y were interleaved with training trials on every session, raising the total number of trials per session from 96 to 100.

##### Experiment 4

Rats received a single phase of training consisting of 42 sessions with two concurrently trained nonlinear discrimination problems. One of these problems was an A(1), X(1), AX(0) negative-patterning discrimination, whereas the other was a B(0), Y(0), BY(1) positive-patterning discrimination. Cues A and B were the same visual cues, and X and Y were the same auditory cues used in the previous experiments, counterbalanced within modality. All trial types were presented 16 times in a session, making a total of 96 trials.

#### Behavioral Measures and Statistical analysis

Conditioned responding was measured in both groups as the number of head entries in the sucrose magazine during the last 5-s of the 10-s cues. Focusing the analysis on the latter half of the cue has two advantages. First, it provides a cleaner measure of goal-tracking behavior, as sign-tracking behavior—which we did not measure, and which may have differed between the groups—tends to concentrate in the first half of a 10-s cue (Holland, 1977). Second, it filters out any bias in conditioned behavior resulting from the fact that Agency and Passive rats began their trials at different locations relative to the sucrose magazine. Indeed, whereas Agency rats necessarily had their snouts in the adjacent nose port at the time of cue onset, Passive rats were free to roam in the chamber and approach the magazine at all times, compromising any between-group comparison at the start of the cue period.

For analysis purposes, the data from each rat was first averaged across trials in a session and further collapsed into average responding across session blocks. Testing and training trial types were analyzed separately due to the high number of trial types and the unequal number of presentations per trial type. A linear mixed-model analysis of variance was used for all data analyses and subsequent post hoc analysis were performed with Bonferroni-corrected significance values. All analyses were conducted in JAMOVI (Gallucci, 2017; The Jamovi Project, 2019).

## Acknowledgements

This research was supported by National Institute on Drug Abuse grants 5R00DA036561 and 1R15DA051795 (GE). We thank Geoffrey Schoenbaum for his comments on a previous version of the MS. The authors declare that they do not have any conflicts of interest (financial or otherwise) related to the data presented in this manuscript. All correspondence to be addressed to Guillem R. Esber.

## Author contributions

Mihwa Kang: Methodology, Software, Investigation, Data curation, Visualization, Formal analysis, Writing-Original draft preparation; Ingrid Reverte: Conceptualization, Validation, Methodology, Software, Investigation, Data curation, Formal analysis; Stephen Volz: Methodology, Formal analysis, Visualization, Writing-Original draft preparation; Keith Kaufman: Data curation, Visualization; Formal analysis; Salvatore Fevole: Data curation, Visualization; Formal analysis; Anna Matarazzo: Data curation, Visualization; Formal analysis; Fahd Alhazmi: Software; Inmaculada Marquez: Software, Visualization; Mihaela D. Iordanova: Conceptualization, Writing-Original draft preparation, Writing-Reviewing and Editing; Guillem R. Esber: Conceptualization, Project Administration, Methodology, Supervision, Writing-Original draft preparation, Writing-Reviewing and Editing, Funding acquisition.

## Author ORCIDs

Guillem R. Esber: https://orcid.org/0000-0001-8874-4255

Mihaela D. Iordanova: https://orcid.org/0000-0001-6232-448X

Ingrid Reverte: https://orcid.org/0000-0002-4120-9324

Inmaculada Marquez: https://orcid.org/0000-0001-7914-6907

## Ethics

Animal experimentation: All animal care and experimental procedures were conducted according to the National Institutes of Health’s *Guide for the Care and the Use of Laboratory Animals*, and approved by the Brooklyn College Institutional Animal Care and Use Committee (Protocol #303).

