## Supplemental Materials for "Agency rescues competition for credit assignment among predictive cues from adverse learning conditions"

**This PDF file includes**

Supplementary tables
Experiments S1, S2

### Supplementary tables

| Exp. | Phase | Total # trials | Mean ITI | SD ITI | Mean session | SD session |
|---|---|---|---|---|---|---|
| 1 | Pretraining | 96 | 18.0 s | 6.7 s | 50. 2 min | 10.7 min |
| | Compound | 96 (100 w/ probes) | 13.1 s | 3.4 s | 43.1 min | 5.6 min |
| 2 | Pretraining | 96 | 19.5 s | 5.8 s | 52.5 min | 9.3 min |
| | Compound | 100 | 18.7 s | 8.5 s | 51.2 min | 13.7 min |
| 3 | Pretraining | 96 | 24.3 s | 7.8 s | 60.1 min | 12.5 min |
| | Compound | 96 (100 w/ probes) | 18.1 s | 7.5 s | 51.3 min | 12.2 min |
| 4 | Patterning | 96 | 13.4 s | 2.7 s | 42.7 min | 4.3 min |
| S2 | Patterning | 96 | 13.0 s | 3.6 s | 42.1 min | 5.8 min |

*Table S1.* Means and standard deviations of the effective intertrial intervals (ITI) and session durations for each experiment and training phase. The effective ITI takes into account not only the programmed ITIs, but also additional between-trial periods introduced by forgone trial offers (trials that were not initiated by Agency rats). Incomplete sessions that timed out after 90 min were excluded from these calculations.

| Exp. | Phase | Session | Agency rat | Passive rat | Completed trials |
|---|---|---|---|---|---|
| 1 | Pretraining | 8 | 5 | 6 | 27/96 |
| | | 2 | 3 | 4 | 72/96 |
| | | 2 | 5 | 6 | 92/96 |
| 2 | Pretraining | 1 | 11 | 12 | 68/96 |
| | | 2 | 11 | 12 | 84/96 |
| | | 1 | 3 | 4 | 84/96 |
| | | 3 | 11 | 12 | 81/100 |
| 3 | Pretraining | 1 | 11 | 12 | 32/96 |
| | | 2 | 11 | 12 | 33/96 |
| | | 3 | 11 | 12 | 42/96 |
| | | 5 | 13 | 14 | 66/96 |
| | | 1 | 15 | 16 | 85/96 |
| | | 1 | 3 | 4 | 89/96 |
| | Compound | 1 | 1 | 2 | 10/96 |
| | | 2 | 1 | 2 | 10/96 |
| | | 1 | 11 | 12 | 74/96 |
| | | 1 | 13 | 14 | 86/96 |
| | | 15 | 11 | 12 | 82/100 |
| | | 14 | 11 | 12 | 90/100 |
| | | 13 | 5 | 6 | 96/100 |
| | | 24 | 15 | 16 | 96/100 |
| | | 13 | 15 | 16 | 96/100 |
| | | 13 | 17 | 18 | 96/100 |
| | | 13 | 7 | 8 | 96/100 |
| | | 13 | 11 | 12 | 96/100 |
| | | 13 | 3 | 4 | 96/100 |
| | | 13 | 13 | 14 | 96/100 |
| | | 31 | 1 | 2 | 99/100 |
| | | 14 | 13 | 14 | 99/100 |
| S2 | Patterning | 25 | 3 | 4 | 24/96 |

*Table S2.* List of sessions that timed out after 90 min before the rats could complete all scheduled trials. Notice that when an Agency rat failed to complete a session, its yoked counterpart in the Passive group was also affected. The ID numbers of all rats concerned are shown for each experiment and training phase, along with the number of trials they completed.

### Exp. S1: Piloting a novel cue-competition task in a standard Pavlovian magazine-approach setting

The purpose of this study was to pilot the novel cue-competition design to be used in the second study in a standard Pavlovian magazine-approach setting. Trials were spaced out by a mean intertrial interval of 60 s. For ease of reference, the experimental design can be found in Fig. S1A.

#### Results

The pretraining phase consisted of 10 sessions of discrimination training with A(1), B(0), X(.75) and Y(.25) (Fig S1B, left). An ANOVA revealed a main effect of cue ($F_{(3)} = 18.77$ , $p < 0.001$) and session block ($F_{(4)} = 16.98$ , $p < 0.001$), but no cue by session block interaction ($F_{(12)} = 1.29$ , $p = 0.231$). Post-hoc analyses of the main effect of cue revealed that responding to A(1) was significantly greater than to B(0) ($t_{(140)} = 5.31$ , $p < 0.001$) and that responding to X(.75) was significantly greater than to Y(.25) ($t_{(140)} = 4.68$ , $p < 0.001$).

The results of the second compound phase, which comprised 20 sessions, are shown in Fig. S1B (right). Competitive cue interactions were evidenced by a gradual switch in responding to X and Y; that is, a decrease in responding to X (due to competition with A) combined with an increase in responding to Y (due to protection from extinction by B). In support of this impression, an ANOVA on responding to X and Y throughout this phase revealed no main effect of cue or session block, but a significant cue by session block interaction ($F_{(9)} = 2.52$; $p = 0.010$). Simple main effects analyses revealed that rats responded significantly more to Y than X on session blocks 9 and 10 ($t_{(140)}$ = [2.44–2.81];

$p = 0.010$). A further ANOVA on the remainder of the cues presented in Phase 2 revealed only a significant main effect of cue ($F_{(3)} = 65.44$; $p < 0.001$). Simple main effects confirmed successful discriminations between A(1) and B(0) ($t_{(280)} = 10.89$; $p < 0.001$) and 3AX(1) and 3BY(0) ($t_{(280)} = 8.25$; $p < 0.001$).

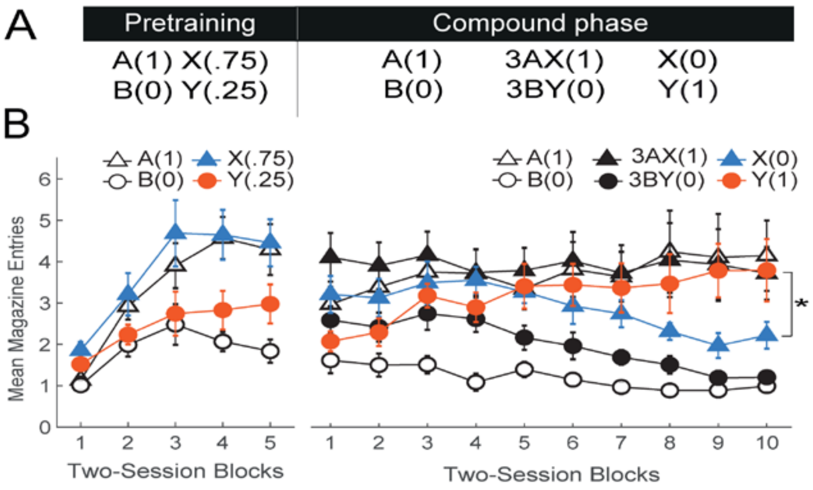


**Figure S1**. Results of a pilot study testing the novel cue-competition design used in Exp. 2 in a standard Pavlovian conditioned magazine-approach setting. Trials were spaced out by a 60-s mean ITI. **(A)** Experimental design. **(B)** Performance during the Pretraining (left) and Compound (right) phases expressed as the mean number of magazine head entries (+/-SEM).

### Materials and Methods

#### Subjects

The subjects were 8 naïve Long-Evans rats. At the start of the experiment they were ~22 wo and weighed 470-527 g (males) and 291-319 g (females). Husbandry details were the same as described in the main text.

#### Apparatus

Same as described in the main text.

#### Procedure

Rats were magazine trained in the manner described in the main text before receiving standard Pavlovian conditioned magazine-approach training. The experimental design was the same as that shown in Fig. 2A. All procedural details were identical to those used in the Passive group of Exp. 2 with two exceptions. First, trials were not preceded by the trial-availability cue (noseport light) and second, the mean ITI was 60 s (range: 30-90s)

### Exp. S2: Ruling out alternatives for the role of agency in competitive credit assignment – Replication

The purpose of Exp. S2 was to replicate the findings of the patterning study in rats that had previously exhibited different degrees of competitive credit assignment. The same Agency and Passive groups used in the blocking task (Exp. 1) went on to receive negative- and positive-patterning problems.

#### Results

Inspection of Fig. S2 suggests that the same Passive rats that showed impaired competitive credit assignment during blocking training were better able to solve these complex non-linear discriminations. A mixed ANOVA on the negative-patterning

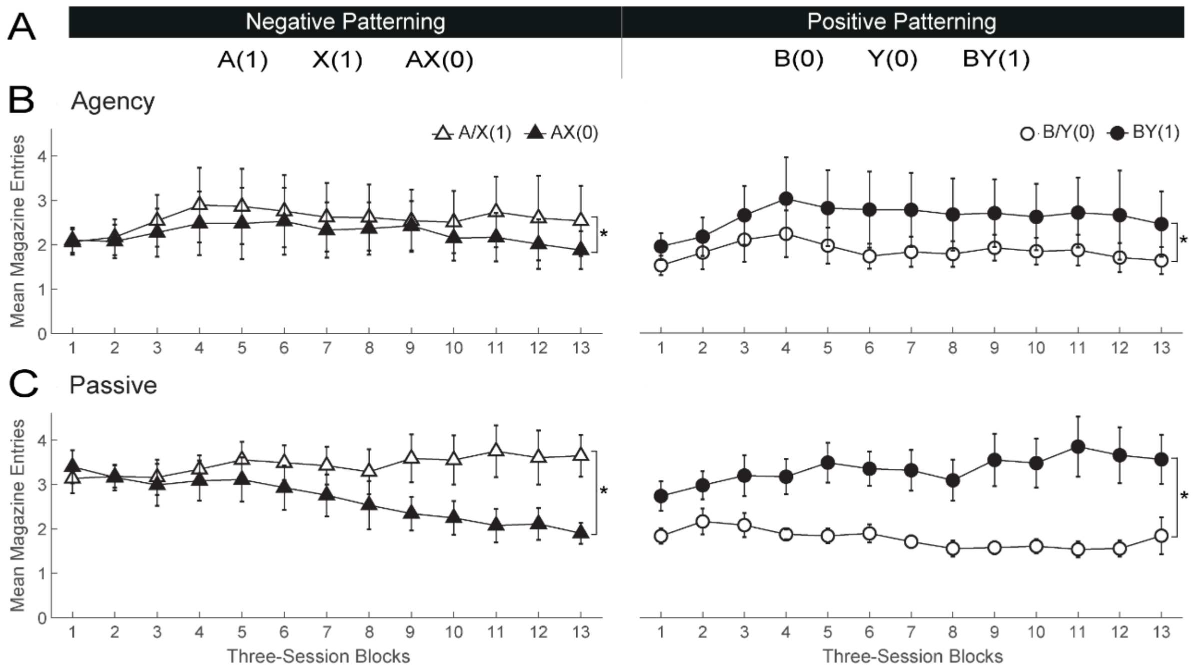


**Figure S2.** Replication of the patterning study (Exp. 4) in rats who had previously received blocking training (Exp. 1). The results confirmed that agency over learning does not rescue competitive credit assignment by enhancing general discrimination proficiency or the ability to process compounded stimuli concurrently.

[A(1)/X(1) vs. AX(0)] discrimination revealed an effect of cue [$F_{(1,350)}$ = 53.37, $p$ < 0.001] as well as significant session block by cue [$F_{(12,350)}$ = 2.31, $p$ = 0.008] and group by cue interactions [$F_{(1,350)}$ = 9.11, $p$ = 0.003]. Simple main effects analysis of the latter interaction revealed that both groups solved this discrimination [Agency: $t_{(350)}$ = -3.03, $p$ = 0.006; Passive: $t_{(350)}$ = -7.30, p < 0.002], although Passive rats showed a larger effect size than Agency rats (Cohen's d = 0.64 and 0.18, respectively). A parallel analysis of the positive-patterning [B(0)/Z(0) vs. BZ(1)] discrimination revealed a main effect of cue (elements vs. compound) [$F_{(1,350)}$ = 172.85, $p$ < 0.001] and a cue by group interaction [$F_{(1,350)}$ = 20.08, $p$ < 0.001]. Simple main effects analyses confirmed that both groups also solved this discrimination [Agency: $t_{(350)}$ = 6.13, p < 0.001; Passive: $t_{(350)}$ = 12.47 , p < 0.001], but, once again, a larger effect was observed in Passive than Agency rats (Cohen's d = 1.50 and 0.48, respectively). These findings bolster the hypothesis that the deficits observed in Passive relative to Agency rats in Exps. 1-3 were specific to cue competition and not the result of impaired processing of compound cues or more general impairments in discrimination learning.

### Materials and Methods

#### Subjects

The subjects were the 16 Long-Evans rats that previously took part in the blocking study (Exp. 1). At the outset of the study they were ~26 wo and weighed 544-601 g (males) and 303-340 g (females). Husbandry details were the same as described in the main text.

#### Apparatus

Same as described in the main text.

**Procedure**

Rats did not require magazine nor shaping training, as they had already received such training at the outset of Exp. 1. The experimental design as well as all other procedures was identical to that used in the patterning study (Exp. 4), with the exception that novel auditory stimuli were used in the role of X and Y (a 2.5-Hz, 80-dB clicker and a 1-kHz, 80-dB tone, counterbalanced). Thirty-nine training sessions were conducted.
